# PDBminer to Find and Annotate Protein Structures for Computational Analysis

**DOI:** 10.1101/2023.05.06.539447

**Authors:** Kristine Degn, Ludovica Beltrame, Matteo Tiberti, Elena Papaleo

## Abstract

Structural bioinformatics and molecular modeling of proteins strongly depend on the protein structure selected for investigation. The choice of protein structure relies on direct application from the Protein Data Bank (PDB), homology- or de-novo modeling. Recent de-novo models, such as AlphaFold2, require little preprocessing and omit the need to navigate the many parameters of choosing an experimentally determined model. Yet, the experimentally determined structure still has much to offer, why it should be of interest to the community to ease the choice of experimentally determined models. We provide an open-source software package, PDBminer, to mine both the AlphaFold Database (AlphaFoldDB) and the PDB based on search criteria set by the user. This tool provides an up-to-date, quality-ranked table of structures applicable for further research. PDBminer provides an overview of the available protein structures to one or more input proteins, parallelizing the runs if multiple cores are specified. The output table reports the coverage of the protein structures aligned to the UniProt sequence, overcoming numbering differences in PDB structures, and providing information regarding model quality, protein complexes, ligands, and nucleotide binding. The PDBminer2coverage and PDBminer2network tools assist in visualizing the results. We suggest that PDBminer can be applied to overcome the tedious task of choosing a PDB structure without losing the wealth of additional information available in the PDB. As developers, we will guarantee the introduction of new functionalities, assistance, training of new contributors, and package maintenance. The package is available at http://github.com/ELELAB/PDBminer.

## Introduction

Structural biology and structural bioinformatics are ever-increasing areas of study. In recent years, structural biology has guided drug discovery [1], shed light on fundamental biology, and aided in biotechnology and bioengineering solutions [2]. Researchers apply structural biology techniques to investigate various facets of protein function and contribute to giving a mechanistic interpretation of biological data. For instance, scientists have employed these methods to study mutations, protein-protein interaction, ligand binding, functional motifs, and alterations [3].

In this context, databases of protein structures, such as the Protein Data Bank (PDB), a community-driven core resource for structural biologists and bioinformaticians [3], are of paramount importance. Structural bioinformatics and molecular modeling have traditionally depended on experimentally solved structures, yet solving protein structures is time-consuming and expensive, and each experimental methodology poses its limitations. As a result, not all proteins are solvable with current techniques, suggesting that fewer protein structures are available compared to the number of discovered protein sequences [4]. Computational structure prediction tools attempt to bridge this gap, relying on available structures in the PDB to train and validate. Since 2021, with the introduction of AlphaFold2 [5] and other tools such as RoseTTAFold [6], the quality of structure prediction has increased tremendously, rendering the predictions significantly more reliable [7]. The subsequent release of the AlphaFold Database (AlphaFoldDB) [4] has remarkably lowered the barrier of entry, allowing many non-computational scientists to take advantage of the predicted structures. While protein structures and predictions for many proteins are freely available, choosing a protein structure for subsequent analysis is not trivial. Some protocols illustrate best practices for electing experimental protein structures [8]. However, they still do not deliver an overview of the available protein structures for a given protein. Searching the PDB database can be a time-consuming task prone to human error due to several common difficulties. These may include inadequate structure coverage of a particular protein in the PDB (e.g., only a domain or a small peptide may be available), the presence of amino acid substitutions or other large-scale sequence variations between the protein sequence in the structure and the UniProt reference, lack of upfront information about protein complex structures involving our target of interest, insufficient information on the presence of other ligands (such as nucleic acids), and differences in numbering between the PDB file and the UniProt sequence.

We present PDBminer, a tool aiding the user in the choice of a protein structure for analysis by mining the PDB and AlphaFoldDB. PDBminer provides the user with information such as the range of amino acids covered by the target protein structure regardless of numbering in the PDB file, quality information about the protein structure itself, details of complexes with other proteins, nucleotides, and ligands, as well as the presence of expression tags.

While many tools exist to handle PDB data, such as PDBrenum [9], a tool integrating AlphaFoldDB and PDB structures, including complex information, alignment coverage, and missing residues does, to the best of our knowledge, not exist.

### Features and Implementation

PDBminer aims to aid the selection of a protein structural model by automating the search in the PDB and the AlphaFoldDB. PDBminer is a Python-based command line tool available at http://github.com/ELELAB/PDBminer under the GNU General Public License v3.0. PDBminer can be used as standalone software or integrated into a pipeline. PDBminer depends on snakemake [10], biopython [11], pandas [12], and biopandas [13] (see the GitHub for versioning details) and on the information in UniProt, AlphaFoldDB, the PDB, and the Protein Data Bank Europe (PDBe) accessed through application programming interfaces (API). The plotting tools require seaborn [14], matplotlib [15], and networkx [16] (see the GitHub for versioning details).

The advantage of PDBminer compared to search on available databases is the automated process. In summary, PDBminer takes the UniProt unique and stable entry identifier (UniProt ID) as input and generates a CSV file that lists all available structures for the protein and their respective details. PDBminer utilizes UniProt, PDBe, and AlphafoldDB APIs to identify PDB structures and relevant metadata, including deposition date, experimental method, and resolution. The found experimental structures are ranked based on their metadata. Each experimental structure sequence is aligned using a pairwise alignment to the reference sequence from UniProt to annotate missing residues, deviations from the UniProt sequence, and mutations. The tool also annotates any other chains or molecules in the structure files. If there are specific mutations of interest, these can be included in the input, and in such a case, the output is filtered to include only structures covering at least one defined mutation site. Additionally, it is possible to define mutational clusters, resulting in filtering based on these clusters. PDBminer reports information deposited in the databases, ready for inspection by the user before using them for further research or processing.

In more detail, the input is, as a minimum, a UniProt ID and a gene name. The output is a CSV file listing all available structures for the UniProt ID and their respective details. PDBminer can be run in two modes, directly using command line options, or employing a configuration file detailing the settings for the run (**figure 1**). Using the configuration file allows the user to specify mutations of interest, mutational clusters, and isoforms. PDBminer will thus filter for structures covering at least one mutation of interest, filter for structures covering a mutational cluster, or align the sequence to the specified isoform.

**Figure 1:**
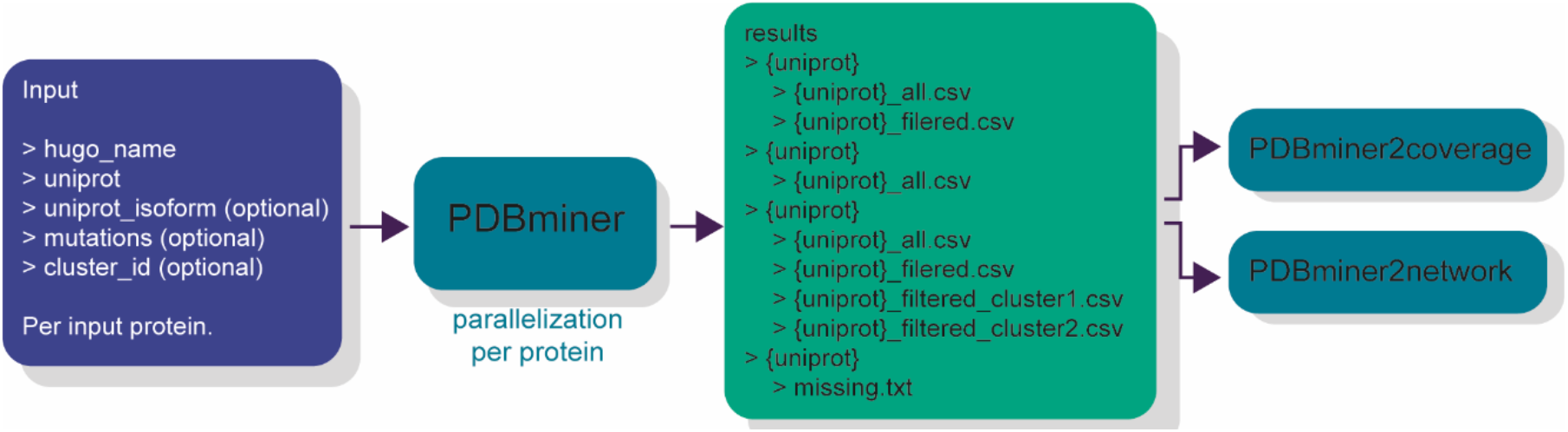
Overview of the Tool. The flow diagram illustrates the input and output of PDBminer. PDBminer takes an input file or command describing as a minimum a gene name (hugo name) and UniProt ID (UniPrsot). sThe output will be either a text file indicating no PDB structures or AlphaFold2 models exist or a {uniprot}_all.csv file detailing the available structures. With more input information, the output becomes increasingly specific. PDB structures are not annotated with their isoform, hence the number refers to the fasta file from UniProt to which the structure sequences are aligned. If mutations are part of the input, the output will be filtered for the models covering at least one mutation. Lastly, it is also possible to annotate clusters of mutations. The input file can be a multi-line file, where the search for models for each protein is parallelized. The plotting modules can be run to visualize the output, which may aid in choosing the appropriate structure.

While just using the command line is convenient for simple searches, providing a configuration file is preferable in two conditions: i) when the objective is to find structures for multiple proteins and one wishes to run them in parallel, ii) when the objective is to find structures covering clusters of mutations. For each UniProt ID in the configuration file or command line, PDBminer uses the UniProt knowledgebase [17] API to identify all PDB structures related to a particular protein, from large oligomeric proteins to small peptides. For each structure, PDBminer initiates a query to the PDBe [18] to assign relevant metadata for each PDB entry identified in the previous step. The metadata includes the deposition date in PDB and the experimental method used to solve the structure, and if the methodology is X-ray crystallography or Cryo-EM, the resolution is also assigned. Since PDBminer utilizes APIs to interrogate UniProt and PDBe, the output of PDBminer depends on the availability of structures in UniProt and PDBe hence, there is a potential lag time. In principle, the deposition of a structure is not reflected immediately in UniProt and thus, not identified by PDBminer. PDBminer identifies the latest AlphaFold2 model available in the AlphaFoldDB, and information corresponding to the metadata is added for the AlphaFold2 model, annotating the experimental method as predicted. The available PDB or AlphaFold2 structures are ranked based on their metadata. The AlphaFold2 model is ranked first, providing grounds for comparison. The PDBs are subsequently ranked following previously published best practices [8]. The ranking utilizes the experimental method in the following order: X-ray crystallography, Cryo-EM, then NMR, followed by other less used methodologies such as Neutron Diffraction and Fiber Diffraction. This information is available in the output file allowing the users to filter based on their needs. There is a ranking within each experimental group based on the resolution for X-ray crystallography and Cryo-EM. If structure resolution is identical, the rank follows their respective deposition dates prioritizing newer structures. Determination of the ranking of entries from NMR and other methodologies follows the deposition date. In cases where a protein has neither experimental structures associated nor an AlphaFold2 prediction in the AlphaFoldDB, a file indicating this is the sole output (**figure 2**).

**Figure 2:**
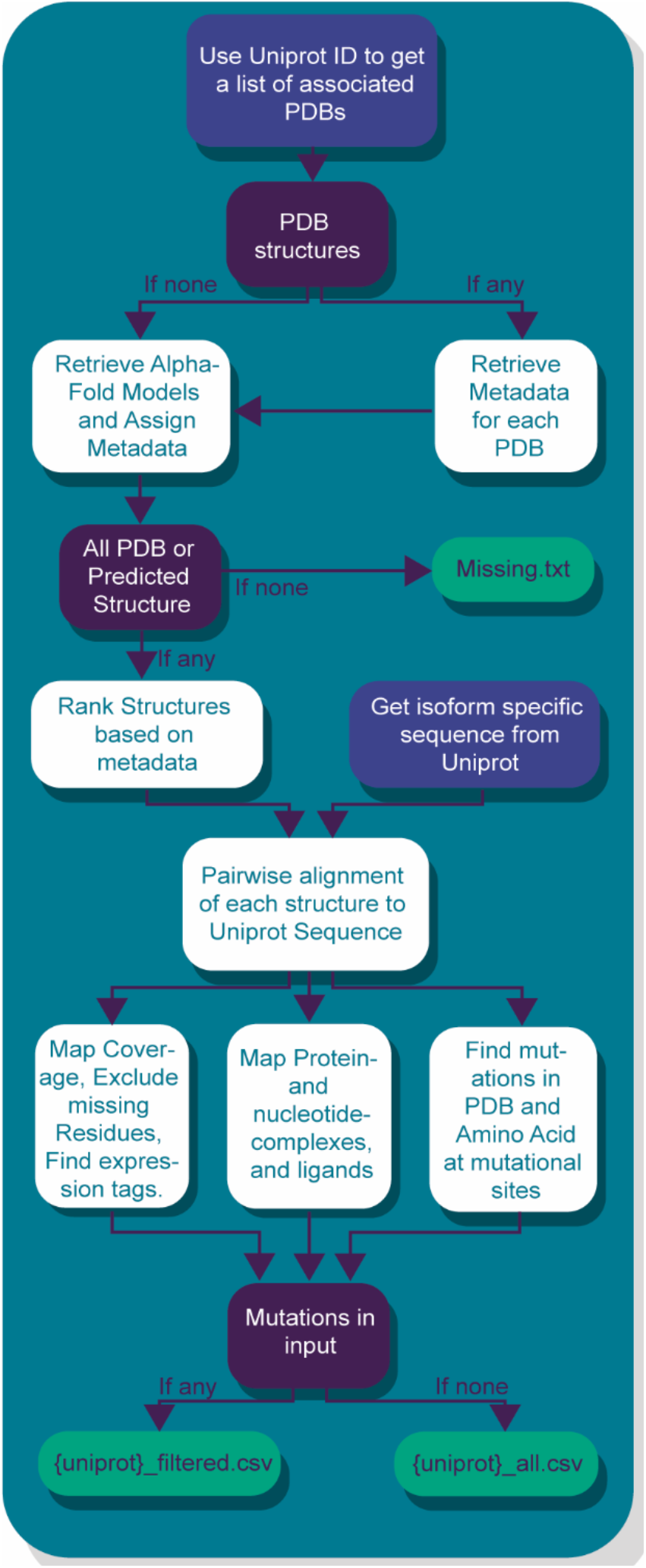
Computational Process of PDBminer. The flow diagram illustrates the PDBminer application architecture. This flow diagram shows how PDBminer uses a UniProt ID to find all related PDB structures with a UniProt API. If any solved structures exist, PDBminer captures the respective metadata (experimental method, deposition date, and X-ray structures resolution) through an API to PDBe. After this, or directly if no PDBs exist, the newest AlphaFold2 model is also retrieved with its relevant metadata, such as version and model date. It ends here when no PDB structures and no AlphaFold2 model exist. If a structure does exist, the workflow continues to the ranking step. The AlphaFold2 structure is ranked highest, then the X-ray crystallographic structures, cryo-EM, NMR, and any other methodology. Each experimental group is ranked to get the best structures near the top. The metadata is available in the output file, allowing the user to re-rank based on the research question. After ranking, PDBminer aligns each model to the appropriate sequence from UniProt. This pairwise alignment allows a renumbering of the PDB structures. PDBminer detects amino acids that differ from the UniProt sequence and identify mutational sites covered by the structure yielding a coverage range per chain, following the UniProt numbering and annotated deviations. For the PDB structures, missing residues will split this coverage into segments. PDBminer uses the same approach for AlphaFold2 models, including residues with a confidence score of 70 or higher. Other elements in the PDB file are mapped, such as protein and nucleotide complexes and lignans or ions. If the input contains mutations or clusters of mutations, PDBminer filters the output to include structures covering at least one input mutation.

Each ranked structure is aligned to the reference UniProt sequence using a pairwise alignment based on a dynamic programming algorithm implemented in the BioPython module to either the specified isoform or the canonical. The goal is to align the often-shorter sequence of the protein structure to the entire UniProt sequence. Accordingly, PDBminer utilizes a local alignment. The alignment setup aims to keep the structure sequence as gapless as possible. Therefore, the alignment rewards a match with 1 point, penalizes a non-match with -10 points, penalizes opening a gap with -20 points, and penalizes a keeping it open with -10 points. The alignment is used exclusively to i) Re-annotate the positions of the amino acids since PDB files often are non-canonically numbered. The alignment score does not constitute an endpoint. ii) To query the sequence for missing residues and other deviations from the canonical UniProt sequence, that is, amino acid substitutions and expression tags. The result of this step is a coverage range indicating the part of the UniProt sequence found in the PDB file, according to the alignment using the UniProt numbering. PDBminer annotates any other chains or molecules, omitting water, available in the structure files. PDBminer annotates if any protein complexes, protein-nucleotide complexes, or other complexes are present. Protein complexes indicate if there are other protein chains in the PDB file and which they are, subcategorized as protein complexes or fusion products. Nucleotide complexes indicate if any DNA or RNA is bound to the protein, and other complexes indicate if any non-protein ligand or metal ion is bound in the structure.

The AlphaFold2 structures undergo a similar alignment process. However, rather than annotating missing residues, any residue with a pLDDT confidence score below 70, meaning those residues whose structure prediction is found to be low confidence, likely due to disorder, is removed from the coverage range. There is no complex query for the AlphaFold2 structures since the current AlphaFoldDB does not contain these.

If the configuration file includes mutations of interest, the output is filtered to include only structures that cover at least one mutation from the list. When using mutation clusters, the filtering relies on each cluster (**figure 2**).

#### Graphic Tools

##### PDBminer2coverage

PDBminer2coverage is a command line tool that generates a plot to provide a visual overview of the structural coverage of the found protein structures generated by PDBminer. The x-axis shows the canonical sequence of the protein, while the y-axis shows the structure models that cover this sequence. **Figure 3A** demonstrates this plot. To access a detailed and true-to-size version visit the GitHub repository.

**Figure 3:**
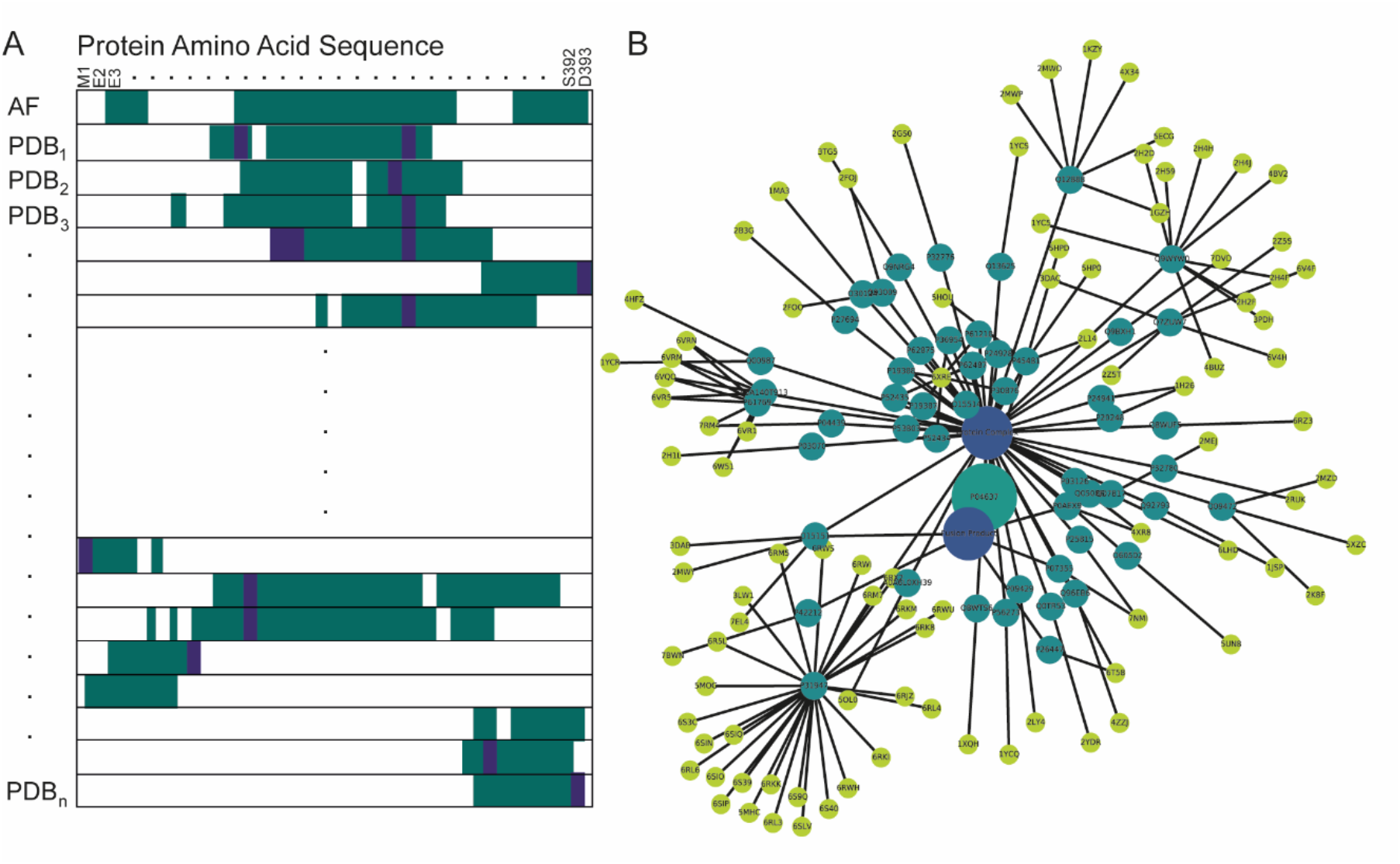
A Case study of TP53. To illustrate the utility of PDBminer, we searched for structures of p53. The search yielded 244 structures. (A) A conceptual drawing of the coverage plot generated for all found structures. Each structure sequence is aligned to the UniProt sequence and colored based on its coverage with teal. The white areas indicate parts of the sequence not covered by the PDB, including missing residues. Mutational sites in the PDB are colored purple. The true-to-size plot is too large to include in a publication but is available in the GitHub repository. (B) A graph showing the protein complexes available in the PDB. This network shows the TP53 UniProt ID P04637 leading to two nodes, Fusion Product and Protein Complex. From each of these are all the found proteins annotated with their UniProt ID, leading to the appropriate PDB identifier.

PDBminer2coverage fills the parts of the UniProt sequence that each structure covers with a solid color, while the areas not covered remain white, including missing residues. When multiple chains in one structure correspond to the protein of interest, the tool annotates each chain with a chain identification letter.

Moreover, any difference in the protein sequence encoded in the PDB file compared to the UniProt sequence is highlighted in a different hue, making it easier to inspect the presence of mutations.

PDBminer2coverage can also visualize mutations of interest used as input in the configuration or command line by overlaying their positions with a transparent secondary color. By default, the parts of the sequence with structural coverage are dark gray, the mutations in the PDB in lighter gray, and the site overlay in blue. Nonetheless, users can choose any color for these inputs using the tool.

##### PDBminer2network

PDBminer2network visualizes the protein complexes found by PDBminer by creating a network graph (**figure 3B**). The network graph plots every bound protein and the associated PDB identifiers as nodes and their internal connection as edges, allowing a visual representation of the protein complexes and fusion products in the PDB.

##### Runtime

The runtime of PDBminer depends on the number of structures that need to be processed. All runs have a short lag time with checks of the API connection and the input formatting. To measure the runtime, we process 12 proteins studied in *MAVISp: Multi-layered Assessment of VarIants by Structure for proteins* [19]. The runtime setup is to i) run each protein individually, ii) run the proteins in a configuration file using one core, iii) two cores, iv) tree cores, and v) four cores. When running the proteins individually, we see the time spent checking API connection and input averages just below a second. The advantage of using the configuration file is that this check only is done once. On average, it takes 2.18 seconds to process each structure. The consistent timing per structure means that processing time is longer for proteins with more associated structures. The time-consuming step in PDBminer is calling the APIs and downloading the PDB files, which are needed to find reliable information on missing residues and realigning. When using one core with the configuration file, the timing is comparable to the command line. When using more cores, runtime decreases **(table 1)**. The tool uses multi-core processing on individual proteins, and the speed of the process is thereby determined by the protein with the highest number of structures. Here the protein with the highest number of structures is KRAS, with 238 PDB files.

**Table 1:**
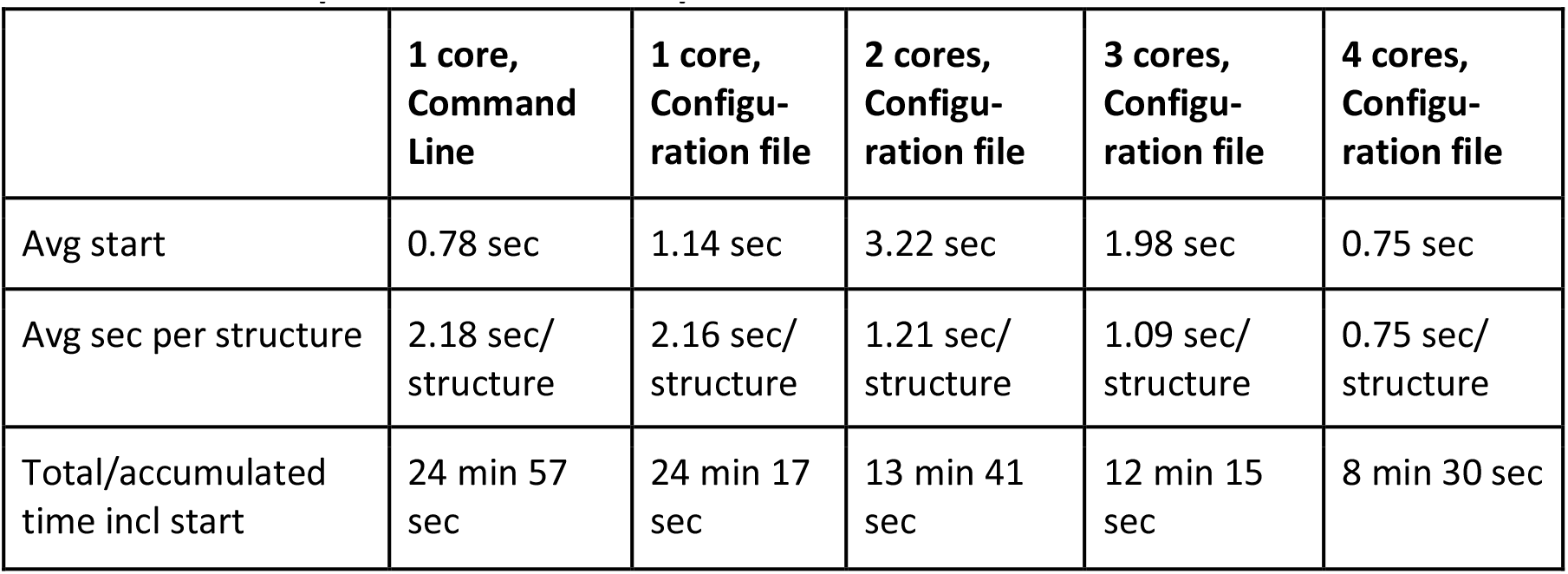
Runtime per structure for all proteins.

|  | 1 core,<br>Command<br>Line | 1 core,<br>Configu-<br>ration file | 2 cores,<br>Configu-<br>ration file | 3 cores,<br>Configu-<br>ration file | 4 cores,<br>Configu-<br>ration file |
| --- | --- | --- | --- | --- | --- |
| Avg start | 0.78 sec | 1.14 sec | 3.22 sec | 1.98 sec | 0.75 sec |
| Avg sec per structure | 2.18 sec/<br>structure | 2.16 sec/<br>structure | 1.21 sec/<br>structure | 1.09 sec/<br>structure | 0.75 sec/<br>structure |
| Total/accumulated<br>time incl start | 24 min 57<br>sec | 24 min 17<br>sec | 13 min 41<br>sec | 12 min 15<br>sec | 8 min 30 sec |

### Case study

We selected TP53 to showcase the results and usage of PDBminer. TP53 is an extensively researched gene due to its significant role as a tumor suppressor gene [20]. Indeed, the p53 protein is involved in cellular functions spanning transcription-dependent and independent processes [21–24]. The protein has included several domains, with examples such as the Zinc binding site (residues Cys176, His179, Cys238, and Cys242), an oligomerization domain (residues at positions 325-356), and the DNA binding domain (residues at positions 91-289). We recently studied the DNA binding domain of p53 with computational methods based on structural biology and predicted the effect of 1132 amino acid substitutions on different aspects of its structure and function. Our investigation required us to collect protein structures in the PDB for the free-state monomer, a dimer, and a DNA-bound version [25]. To this extent, we used PDBminer to gain an overview of the available structures. This overview allowed us to shortlist protein structures from the vast space of p53 structures that satisfied our needs without limiting ourselves to structures previously used for similar studies with different endpoints or spending monotonous time manually and potentially unstructured curating the databases. Hence, the benefit of using PDBminer is to summarize the PDB while earnestly peeling away any mismatch in numbering between models, differences in the sequence such as missing residues and mutations, and annotating all other elements in the PDB file, thereby creating a complete multifaceted overview of the available structures.

Here, we ran PDBminer on p53 from the command line with the gene name TP53 and the UniProt ID P04637. This run resulted in the output file P04637_all.csv containing 244 proteins and peptides covering parts of the p53 protein. Below are four examples of use cases and the subsequent choice of structure aided by the file P04637_all.csv.

#### 1. Study the Free Wild-Type of the DNA binding domain

We utilized PDBminer to find a structural model covering the DNA binding domain. The goal was to study a wild-type structure in a free state. Thus, we consulted the coverage plot and the output file, P04637_all.csv (demonstrative in plot **figure 3A**. To access a detailed and true-to-size version visit the GitHub repository). The output file contained the ranking of the protein structures, mirrored in the coverage plot. Hence, upon visual inspection, we saw how the top-ranked structures included missing residues and mutations. Since the goal was to study the wild-type, we discarded these structures, inadvertently illustrating how the models with the finest resolution can have other limitations. We filtered the output file to remove structures in protein and nucleotide complexes aiming to retain only free structures. Consequently, we had just 18 PDB entries out of the initial 244. The two structures ranking highest after the filtering were the AlphaFold2 model (AF-P04637-F1-model_v4) and 2XWR (1.68 Å) [26]. 2XWR is a zinc-bound X-ray structure solved in 2010. Either of them could have been a good starting point for our investigation. The coverage plot and output file provided three tangible improvements to a search in the PDB: i) The alignment within PDBminer allowed us to compare the coverage of different structures, ii) mutations and missing residues were clearly on display, which is often not the case in PDB entries and iii) we could filter based on binding partners.

#### 2. Study the conformation in the DNA binding

To investigate the conformational changes upon DNA binding and the effect of mutations on DNA binding, we first used PDBminer to identify a suitable starting structure of the DNA-bound p53 DNA binding domain. We used the CSV output file to filter for nucleotide complexes. Thereby, we found 40 structures bound exclusively to DNA or RNA. In 21 cases, the sequence of p53 is unchanged compared to the UniProt sequence, while the remaining 19 harbor mutations. Here, we wished to have a wild-type structure. Thus, the highest-ranked models satisfying these criteria were 7B4N (resolution of 1.32 Å) and 7B4H (resolution of 1.39 Å) [27], which are X-ray crystallography structures from 2020 solved to study oncogenic reactivation of p53. A drawback of both is that they contain missing coordinates on multiple residues, which may require reconstructing, for example, by using MODELLER [28]. While this could be a suitable solution, in this setup we discarded these experimentally solved structures. The next set of models in the ranking was 5MCT (1.45 Å), 5MG7 (1.45 Å), 5MF7 (1.59 Å), and 5MCV (1.6 Å) [29], which are X-ray structures with no mutations, solved in 2016 to study the impact of the DNA conformation on binding. The DNA of 5MCT and 5MCV is a synthetic construct that may or may not be fit for a particular downstream study. Hence, we also discarded these models. Thus, reviewing the 19 structures, aiming to find one without missing residues and with natural DNA, the conclusion is to use either 2AHI (1.85 Å) or 4HJE (1.91 Å). 2AHI is an X-ray crystallography structure from 2005 [30] solved to study the structure of p53 upon DNA binding. 2AHI contains two sets of dimers binding discontinuous DNA. 2AHI is a fine choice to study single chain or dimer binding of DNA. The alternative, 4HJE, is an X-ray crystallography structure from 2012 [31]. 4HJE is a tetramer and is a good choice when studying tetrameric DNA binding. This use case illustrates how choosing a structure depends on the research question of interest. In this case, PDBminer provided the same tangible improvements as the first use case, here with a different purpose on the binding partner parameter.

#### 3. Study the conformational change or protein interaction for mutations on specific positions

One of the central investigations for computational biologists is to study the effects of mutations at sites. To this effect, it is relevant to explore the presence of the mutant in an existing PDB entry. The purpose of finding such a model is twofold. To give an idea of conformational changes and to use the mutant in comparison with a wild-type. An example of this is residue R249 located near the DNA binding site. R249 is a hotspot site for cancer mutations [32] and researchers have shown how the R249S variant has lost the capacity to activate the Bax/Bak complex and thus mechanistically inhibit apoptosis, hence losing one of the main effects of p53 [33].

To find structures with mutated R249, we consult P04637_all.csv, filtering for the presence of mutations on the site. This filtering resulted in 12 potential models where the top three ranked structures are 3D06 (1.2 Å) from 2008 harboring R249S, 3D08 (1.4 Å) from 2008, or 3IGK (1.7 Å) from 2009 carrying both H168R and R249S. 3D06 and 3D08 were solved to study repressor mutations [34], and 3IGK was solved to investigate diversity in DNA binding [35]. In this case, the choice of structure is 3D06, allowing downstream analysis of the variant.

#### 4. Study of Protein-Protein interactors, overview, and binding of HPV virus

In another study design, we want to map the interaction sites and study protein complexes of p53. To gain an overview of solved complexes, we consult the network generated by PDBminer2network (**Figure 3B**). This plot illustrates the proteins experimentally solved in complex or fusion with p53. In the case of p53, PDBminer found 50 different proteins in 89 PDB files. For some complexes, the PDB contains numerous structures. An example of this is SNF (UniProt: P31947), a G2/M progression inhibitor regulated by p53 [36]. The PDB contains 28 experimentally determined structures of the p53-SNF complex. Contrarily, some complexes are only present in one PDB file, such as RPA1 (P27694), a protein involved in DNA metabolism [37,38].

As an example of a protein complex of interest, we identified 4XR8 (2.3 Å) from 2015 covering HPV16 E6/E6AP/p53 ternary complex [39] solved to study how the viral oncoprotein E6 binds and subsequently degrades p53. This viral hijacking process takes part in viral-induced cancers. Investigating the binding can provide a deeper understanding of the process and why this structure could be a starting point for studying interactions between p53 and E6, for example, by applying protein structure networks such as PyInteraph [40].

Collectively PDBminer can aid the selection of structure for research by mining the PDB and may offer justification to choose or opt out of using a particular protein structure.

## Conclusion

Choosing the starting structure of a protein for computational structural analysis is not a trivial task. Indeed, searching through the PDB can be daunting due to the sheer volume of available data. We presented PDBminer, a tool to overcome common challenges in choosing a protein structure for computational research. The advantages of PDBminer include the overview of available structure models and their realignment to the UniProt sequence, omitting challenges with numbering, the outline of mutations within the PDB files which is not readily available information, the option to filter based on specific sites allowing a restricted output, the annotation of binding partners that offers a way to filter based on research needs and finally the flexible graphic tools that makes visual inspection easy. One core use for PDBminer is in the high-throughput application of structural methods for disease-related variant classifications [19]. A tool such as PDBminer is necessary to streamline the selection of the structure of proteins and their complexes. In conclusion, PDBminer allows the user to make an informed choice for further studies.

## Data and Software Availability

The data presented in this paper is publicly available via UniProt and the PDB. The software is available at http://github.com/ELELAB/PDBminer. Here you will also find details to reproduce the case studies presented here.

